# Predicting gene level sensitivity to JAK-STAT signaling perturbation using a mechanistic-to-machine learning framework

**DOI:** 10.1101/2023.05.19.541151

**Authors:** Neha Cheemalavagu, Karsen E. Shoger, Yuqi M. Cao, Brandon A. Michalides, Samuel A. Botta, James R. Faeder, Rachel A. Gottschalk

## Abstract

The JAK-STAT pathway integrates complex cytokine signals via a limited number of molecular components, inspiring numerous efforts to clarify the diversity and specificity of STAT transcription factor function. We developed a computational workflow to make global cytokine-induced gene predictions from STAT phosphorylation dynamics, modeling macrophage responses to IL-6 and IL-10, which signal through common STATs, but with distinct temporal dynamics and contrasting functions. Our mechanistic-to-machine learning model identified select cytokine-induced gene sets associated with late pSTAT3 timeframes and a preferential pSTAT1 reduction upon JAK2 inhibition. We predicted and validated the impact of JAK2 inhibition on gene expression, identifying dynamically regulated genes that were sensitive or insensitive to JAK2 variation. Thus, we successfully linked STAT signaling dynamics to gene expression to support future efforts targeting pathology-associated STAT-driven gene sets. This serves as a first step in developing multi-level prediction models to understand and perturb gene expression outputs from signaling systems.

## Introduction

Cells interpret extracellular cues through biochemical signaling pathways that regulate the activity and specificity of transcription factors (TFs) to control gene expression [1]. Janus kinases (JAKs) and signal transducers and activators of transcription (STATs) constitute a signaling-to-transcription network that operates downstream of more than 50 cytokines and growth factors to induce diverse, yet context-specific gene expression and functions, including the regulation of inflammation, differentiation, and cell survival [2–5]. As in many other well-studied signaling systems, there is a detailed understanding of the regulatory mechanisms that shape cytokine-induced signaling dynamics, a growing understanding of the diversity of STAT-mediated transcriptional responses, but a limited ability to link signaling dynamics to downstream gene expression. The ability to predict signaling features that lead to the expression of specific genes would greatly facilitate rational targeting of cellular function for therapeutic benefits.

The JAK-STAT pathway has long been studied in the context of functional diversity because a limited number of JAK and STAT molecules integrate a broad range of cytokines to elicit varying patterns of gene expression [2–5]; the combinatorial complexity of receptor-JAK-STAT interactions is not sufficient to distinguish context-specific functional responses. Numerous potential mechanisms for achieving STAT diversity and specificity have been interrogated, including variation in post-translational modifications impacting STAT-DNA binding [4], the relative downstream activity of different STAT dimers [6], and the cooperation with additional TFs, such as the interaction of STATs and IRFs to support non-canonical STAT DNA binding and gene transcription [7, 8]. Here we focus on the temporal coding theory, which postulates that stimuli are encoded by the temporal dynamics of TF activation, by measuring STAT-activating tyrosine phosphorylation (STAT3 Y705 and STAT1 Y701) that induces dimerization, nuclear localization, and DNA binding [9]. We hypothesize that STAT tyrosine phosphorylation dynamics are associated with additional unmeasured signaling events that impact the ability of STATs to interact with downstream binding partners to support the transcription of distinct gene sets.

Temporal coding has been extensively investigated in the NF-kB system, in which dynamic features of TF activation have been linked to stimulus-specificity in some studies [10] and select gene induction in others [11]. In the JAK-STAT system, previous studies have noted that STAT-activating tyrosine phosphorylation sites exhibit unique dynamic patterns depending on the stimulating cytokine and that various negative regulators play a role in those cytokine-specific temporal behaviors [12–14]. For example, suppressor of cytokine signaling 3 (SOCS3) shapes cytokine-specific STAT dynamics and shifts the profile of key inflammatory genes from pro-to anti-inflammatory in specific macrophage contexts [12, 14, 15], supporting the connection between signaling dynamics and downstream gene induction. Despite our knowledge that cytokine response diversity exists at both the signaling and transcriptional level, the critical links between differential TF activation and gene expression are unclear. Up to now, studies have relied on modulating cytokine-induced STAT activity experimentally via inhibitors or knockouts and measuring select gene outputs to gain insight into the overall cellular response [13]. However, given the diversity of cytokine-induced gene expression, a more complete understanding of how pSTAT dynamics impact cell function, requires investigation of how dynamics are disparately decoded by genes across the transcriptome.

Inferring dynamic links between signaling events and transcriptomes requires a systems approach. Previous studies have utilized mechanistic modeling to systematically evaluate network components and architecture supporting JAK-STAT signaling dynamics [13, 16], while others have generated multi-omics datasets to understand how targeted JAK-STAT pathway alterations impact gene expression and functional outcomes [16, 17]. While both types of approaches have yielded key insights into cytokine-specific JAK-STAT signaling and gene expression, none have taken an agnostic approach to understanding signaling mechanisms and their links to global gene expression. In fact, across diverse signaling systems, there are few examples of approaches that link signaling to global gene expression [18]. Mechanistic signaling models are generally adept at recapitulating a few key dynamic features from experimental data and can identify points of regulation that mediate responses of interest. However, these models lack the scalability to predict extensive biological outcomes, like global gene expression, due to the vast number of reactions necessary to reproduce such behaviors, and due to the sheer amount of experimental data needed to constrain such a model. On the other hand, more data-driven approaches, including most machine learning models, are capable of wide-scale predictions given enough data, but are difficult to extract biologically meaningful insights from, due to the lack of causal relationships inferred from the data.

Balancing the strengths and weaknesses of individual modeling approaches, we have created an integrated mechanistic-to-machine learning model to analyze signaling mechanisms that shape STAT phosphorylation dynamics and identify genes predicted by STAT signaling dynamics. We model IL-6 and IL-10-induced JAK-STAT signaling in macrophages, key immune cells that integrate cytokines and other extracellular cues to promote or resolve inflammation [19]. IL-6 and IL-10 signal through common STATs (STAT1, STAT3) with unique transcription factor dynamic patterns [13] and induce diverse, but distinct, gene sets associated with contrasting inflammatory functions [15, 20]. Thus, this is an optimal model system for interrogating context-specific JAK-STAT activity.

With our mechanistic-to-machine learning JAK-STAT model, we recapitulated experimental data of IL-6 and IL-10-induced STAT phosphorylation, including well-described cytokine-specific features, and predicted cytokine-specific gene sets associated with late STAT3 dynamics using data-driven machine learning as a means to validate our pipeline. Towards a more quantitative understanding of this signaling network, we performed an in-depth model parameter analysis using our mechanistic model and identified key signaling relationships supporting the regulation of STAT dynamics. Finally, we perturbed the identified signaling components within the mechanistic model to make predictions at the levels of pSTAT dynamics and global gene expression. This work represents an important proof of concept, demonstrating multi-level computational predictions upon signaling perturbations. This workflow is generalizable to other signaling-to-transcription networks and may support future efforts to therapeutically target desired sets of genes and cellular function via the associated signaling features.

## Results

### A mechanistic-to-machine learning modeling workflow for multi-level signaling-to-gene predictions

We have developed a mechanistic-to-machine learning model workflow that leverages the strengths of multiple modeling modalities to make complex multi-level signaling predictions (Fig. 1A). In this workflow, a literature-based mechanistic model of cytokine-induced JAK-STAT signaling is fit to timeseries STAT phosphorylation data, producing large sets of simulated phospho-STAT trajectories. These model trajectories then serve as input to a supervised machine learning model that relates cytokine-induced trajectories to corresponding cytokine-induced gene expression profiles obtained from RNAseq data. We used mechanistic model trajectories for the gene expression predictions, as opposed to experimental signaling data, because this allowed us predict changes in global gene expression in response to perturbation of signaling prior to the generation of additional datasets.

**Figure 1.**
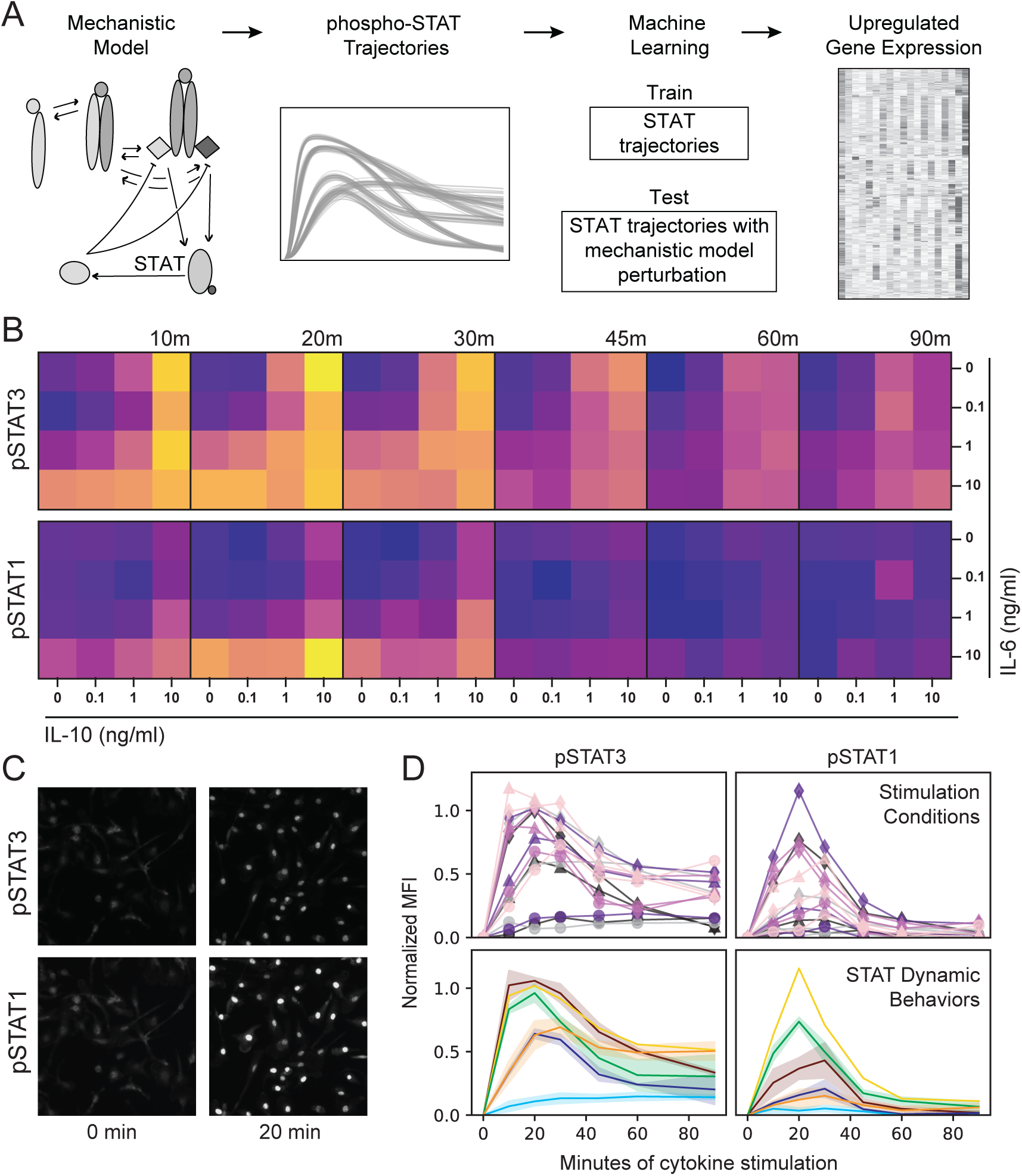
Varied IL-6 and IL-10-induced STAT phosphorylation dynamics support an integrated mechanistic-to-machine learning modeling workflow. (A) Integrated modeling workflow in which mechanistic model simulated pSTAT trajectories are utilized for the training and testing of a machine learning model predictive of transcriptomic data. (B-D) BMDM were stimulated with varying concentrations of the cytokines IL-6 and IL-10, alone and in combination, and STAT3 (Y705) and STAT1 (Y701) phosphorylation was quantified by IF. (B) Mean fluorescence intensity (MFI) was averaged across 3 representative experiments (dark = min MFI, light = max MFI). (C) Representative images of IL-6 10 ng/ml stimulation. (D) IL-6 and IL-10-induced pSTAT1 and pSTAT3 from the 15 stimulation conditions shown in (B) were averaged across all experiments and grouped into 6 behaviors with K-means clustering.

Our approach required linked pSTAT and RNAseq datasets in the context of IL-6 and IL-10 stimulation conditions that maximize diversity in STAT3 and STAT1 activation dynamics. To identify optimal conditions, we stimulated bone marrow derived macrophages (BMDM) with 15 unique dose combinations of IL-6 and IL-10 for up to 90 minutes and measured pSTAT3 (Y705) and pSTAT1 (Y701) using high-content immunofluorescence (IF) imaging (Fig. 1B-D). Consistent with the existing literature [3, 12, 13], (1) both cytokines induced robust pSTAT3 responses, with more sustained phosphorylation in response to IL-10 than IL-6, and (2) IL-6 induced stronger pSTAT1 responses compared to IL-10 (Fig. 1B, Fig. S1A). Although high content IF visualizes pSTAT in single cells, we found that Mean Fluorescence Intensity (MFI) was reflective of the population response over time (Fig. S1B), and so chose to use MFI as the readout for our pipeline. We used K-means clustering to group cytokine conditions resulting in similar phosphorylation dynamics across both STATs and defined six patterns of behavior that the cells displayed in response to IL-6 and IL-10 (Fig. 1D). We selected six stimulation conditions reflecting these STAT behavior patterns (Fig. S1A), with the exception of the non-responding group (light blue), to use in the signaling-to-gene expression modeling workflow.

### Rule-based mechanistic model of JAK/STAT signaling recapitulates diversity in IL-6 and IL-10 induced STAT1 and STAT3 phosphorylation dynamics

We developed a rule-based mechanistic model of IL-6 and IL-10-induced JAK-STAT signaling in BMDMs, in which molecules are represented as structured objects and biochemical interactions between them are described by reaction rules [21]. For example, IL-6 and IL-6R are molecules, with the latter containing a ligand-specific binding site, and binding between these molecules is described by a reaction rule that reversibly creates a bond between these molecules (more detailed description in Methods). The literature-based model of IL-6-induced or IL-10-induced STAT1 and STAT3 phosphorylation consists of 31 reactions rules and 46 unknown model parameters (e.g., kinetic rates, protein concentrations; Fig. 2A, Table S1). In brief, IL-6 and IL-10 utilize separate receptors and go on to phosphorylate STAT1 and STAT3 downstream through the relevant JAK proteins. We have modeled JAKs interacting with both the IL-6 and IL-10 receptors (TYK2 and JAK1) as JAK1, and JAK2 with only IL-6 based on literature evidence [22–24]. Negative regulators, including transcriptionally induced SOCS proteins and the constitutively expressed protein tyrosine phosphatases, PTP1 and PTP3, are included to modulate the dynamic response.

**Figure 2.**
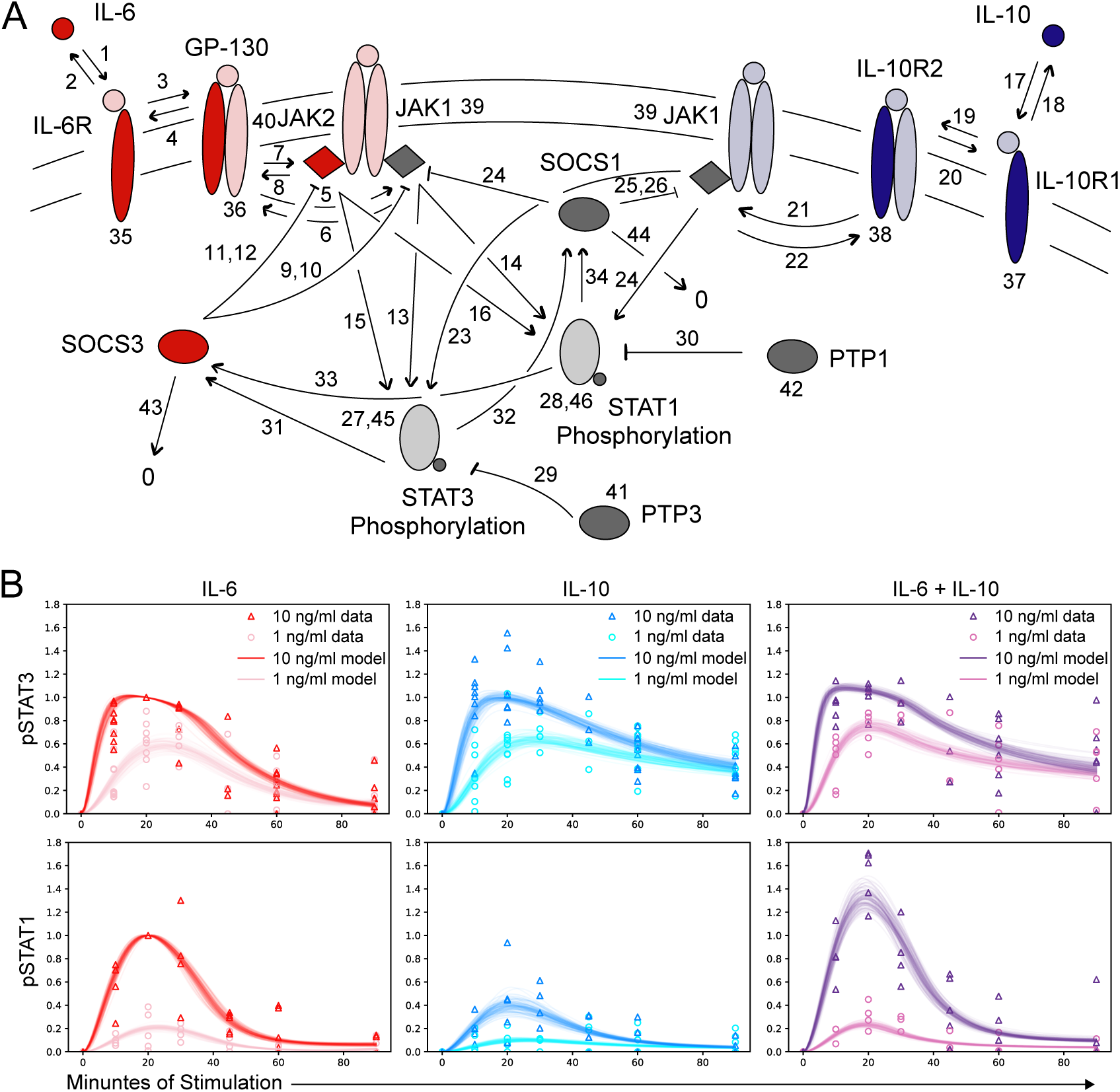
Rule-based mechanistic model of JAK/STAT signaling recapitulates diversity in IL-6 and IL-10 induced STAT1 and STAT3 phosphorylation dynamics. (A) Model architecture including cytokines (IL-6, IL-10), receptor components (IL-6R, GP130, IL-10R1, IL-10R2), kinases (JAK1, JAK2), transcription factors (STAT1, STAT3), induced negative regulators (SOCS1, SOCS3), and protein tyrosine phosphatases (PTP1, PTP3). Numbers represent unknown kinetic parameters, colored components note model proteins (red = IL-6-specific, blue = IL-10-specific, gray = shared), and dark colored components denote the specific reaction occurring according to defined reaction rules (see methods). (B) Cytokine-induced pSTAT timecourse data from IF (points represent independent experiments normalized to IL-6 10ng/ml at 20 minutes) and model fits from approximately 2000 unique parameter sets (curves; see methods for parameter estimation).

The unknown parameters were estimated using an iterative model fitting process that generates an ensemble of parameter sets, each of which generates a model trajectory that matches experimental data points with comparable accuracy [25]. During the fitting process, the associated model parameter values become more constrained, ideally approaching a log-normal distribution (Fig. S1C; more detailed description in Methods). After completing the parameter estimation, we obtained approximately 2000 parameter sets producing model-simulated pSTAT trajectories that recapitulated key features of experimental data, including cytokine-specific pSTAT3 duration and pSTAT1 amplitude (Fig. 2B).

### Modeling workflow identifies late STAT3 activity as predictive of cytokine-specific gene expression

pSTAT3 responses have been well-described in the literature to display cytokine-specific dynamics [13], but the unique signaling features have not been systematically linked to gene expression. To validate the capabilities of our workflow to predict global gene expression from mechanistic model generated trajectories (Fig. 1A), we sought to identify gene sets predicted by the early versus the late pSTAT3 dynamic response (Fig. 3A). This is an optimal test case for our workflow because IL-6 and IL-10-induced pSTAT3 dynamics are comparable in the early response but differ significantly at later time points (Fig. 2B). Thus, we expected that late pSTAT3 would be more predictive of cytokine-specific gene clusters than early pSTAT3. We simulated pSTAT3 trajectories across stimulation conditions and used an unbiased method (Fig. S2) to identify timeframes corresponding to the early (0-13 minutes) and late (50-90 minutes) cytokine-induced pSTAT3 response (Fig. 3B). These simulated trajectory timeframes were used to predict gene expression quantified in response to the same six stimulation conditions; BMDM were stimulated with two concentrations (1 ng/ml and 10 ng/ml) of IL-6, IL-10, alone and in combination, for 1, 2, and 4 hours, resulting in 1592 genes that were upregulated by a minimum of 1.5-fold in at least one condition (Fig. 3C; left).

**Figure 3.**
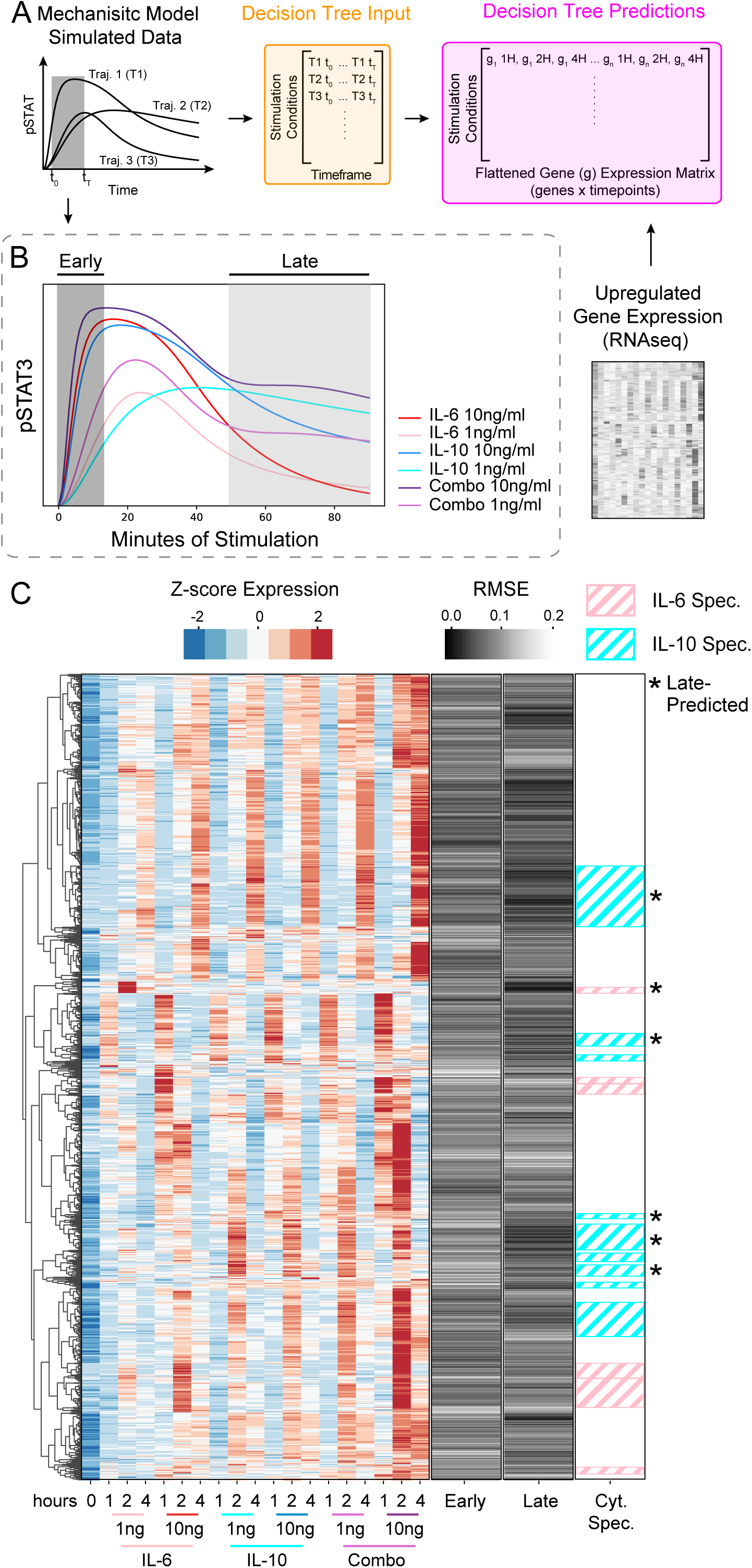
Modeling workflow identifies late STAT3 activity as predictive of cytokine-specific gene expression. (A) Detailed Decision Tree Regressor prediction process using mechanistic model generated pSTAT trajectories. Biologically relevant portions of the trajectory (early and late timeframe) were used as input to the Decision Tree Regressor and flattened, normalized gene expression values were used as labels. (B) Early and late timeframes of pSTAT3 trajectories used for gene predictions. For example, the early timeframe from the IL-6 1 ng/ml trajectory would be the input for the machine learning model, and the output would be the corresponding gene expression values across time for IL-6 1 ng/ml. (C) Hierarchical clustering of upregulated, z-score normalized DEGs across pooled samples (n = 2), gene expression prediction errors (RMSE) using the early versus late timeframes, and cytokine-specific gene clusters with at least 10 genes per cluster. Pink diagonal groups are IL-6-specific gene clusters, while cyan diagonal groups are IL-10-specific clusters. Asterisks denote cytokine-specific clusters that were well-predicted by the late timeframe (see methods).

To identify gene sets associated with the different portions of the pSTAT3 response, we constructed decision tree regressor models for the early and late response, separately. Each model used the appropriate portion of the pSTAT3 trajectory to construct a decision tree regressor model to predict gene expression as a function of the pSTAT3 dynamics (Fig. 3A). The output predictions of the machine learning model were the flattened (across time) gene expression values from our RNAseq data under the various stimulation conditions. Gene predictions were evaluated against the acquired RNAseq expression values using the root mean squared error (RMSE), where lower values represent more accurate gene predictions while larger values represent less accurate predictions. RMSEs were averaged across each time point to generate gene predictions for the early versus late pSTAT3 profiles (Fig. 3C; middle two bars). The early and late timeframes exhibited different patterns of gene predictions, with the early timeframe having a range of RMSEs between 0.0215 and 0.2436, and the late timeframe values ranging from 0.0079 to 0.2042.

To test our hypothesis that cytokine-specific genes would be better predicted by the late pSTAT3 response, we aimed to identify cytokine-specific gene clusters from the RNAseq data (Fig. 3C; left). We used hierarchical clustering to identify 71 total gene clusters, out of which 15 were defined as cytokine-specific (Fig. 3C; right bar; see methods). Cytokine-specific gene cluster predictions were obtained by averaging the RMSEs of genes within the cluster, and the clusters with average RMSEs in the lowest 1/3 were considered relatively well-predicted. This procedure was conducted for both the early and late timeframe predictions and identified six cytokine-specific gene clusters as late-predicted (Fig. 3C; right bar), while no cytokine-specific gene clusters were relatively well predicted by the early response. Thus, the stronger association of cytokine-specific genes with the late pSTAT3 response, compared to the early response, is consistent with the literature and our data demonstrating that cytokine-specific STAT phosphorylation patterns arise at later time points. This gives us confidence that our computational workflow is linking signaling to gene expression in a biologically meaningful way and provides some insight into the specific gene sets that are associated with the late pSTAT3 response (see supplementary file for gene lists).

### Correlation analysis of unconstrained mechanistic model parameters uncovers interpretable signaling relationships

The goal of our integrated framework is to enable identification of signaling components that regulate STAT phosphorylation and to predict the effect of targeted signal perturbations on global gene expression. Interpreting the impact of quantitative variation in model parameters (e.g., protein expression values or activation rates) first requires that model parameters are constrained by the available experimental data. Unconstrained models can be difficult to interpret because effects of variation in individual model parameters on signaling dynamics can be obscured by coincident changes in different, albeit related parameters (e.g., binding and unbinding rates). Thus, we developed a methodology to identify groups of related parameters that are together constrained by the existing data, rather than individually, to support further model analysis. In our ensemble of fit parameter sets, we found that only five of our 46 parameters were well-constrained (log-normally distributed and values within a 2-log range) (Fig. S3A). We then calculated correlations across parameter sets and used hierarchical clustering to group together strongly correlated parameters (Fig. 4A). While some related parameters were expected, such as SOCS1_jak1 binding and unbinding, many identified parameter relationships were less obvious. For example, the top left group of correlated parameters (Fig. 4A) includes IL10-induced JAK1-mediated STAT3 activation and STAT1 protein levels (Fig. 4A); while we appreciate the complexity of molecular interactions, some of these relationships are hard to deduce from biological intuition alone.

**Figure 4.**
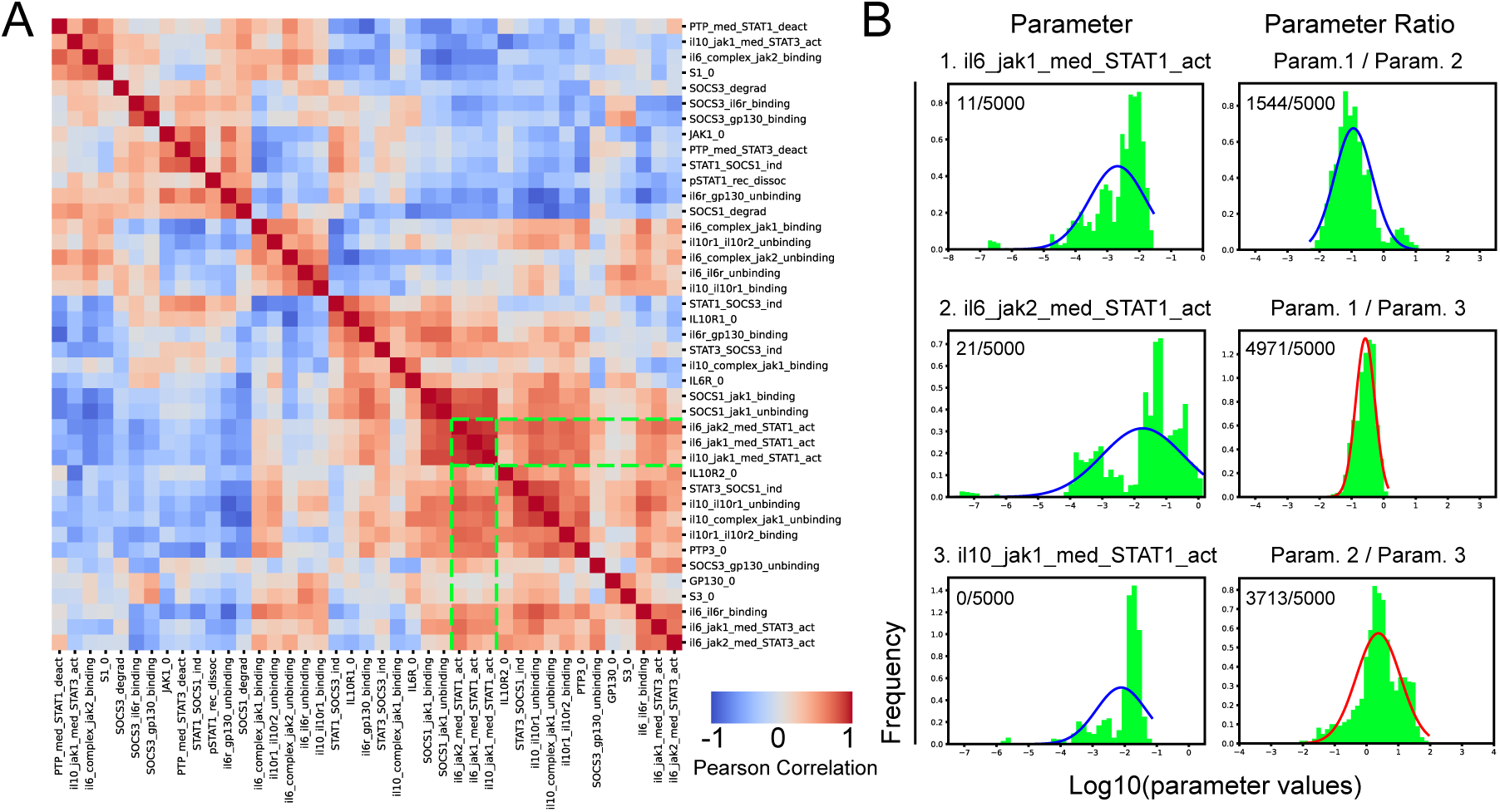
Correlation analysis of unconstrained mechanistic model parameters uncovers interpretable signaling relationships. (A) Hierarchical clustering of Log_10_ parameter correlations across all data fit parameter sets. Colors represent correlation values (red = positive, blue = negative) and green dotted lines note parameters analyzed in B. (B) Distributions of individual parameters (left column; Param 1, 2, and 3 from top to bottom) and parameter ratios (right column). Shapiro-Wilk normality testing results denoted in corners, X out of 5000 bootstrapped samples, with red curves showing distributions deemed log normally distributed in more than half of the bootstrapped runs.

While unexpected parameter relationships are worth exploring further, we were intrigued by a more intuitive, highly correlated group of JAK related parameters (Fig. 4A, green lines). Given the number of selective JAK inhibitors on the market that aim to modulate cytokine-specific function in various inflammatory conditions with undesired off-target effects [3, 26, 27], we were particularly interested in exploring JAK-associated model parameters that could provide insights into JAK-STAT relationships that regulate specialized gene sets. Our parameter analysis identified three correlated parameters describing the ability of JAKs to activate STAT1 that, individually, were poorly constrained by the experimental data (Fig. 4B; left column), but when combined as ratios became well-constrained ratio distributions (Fig. 4B; right column). In contrast, random, chosen parameter pairs did not have constrained ratio distributions (Fig. S3B).

Given well-constrained ratios of JAK-to-STAT1 activation rate parameters, we could then make predictions about how relative changes in these parameter values impact STAT dynamics and gene expression. We found that the logged ratio of JAK1-STAT1 activation and JAK2-STAT1 activation were mostly below 0 (Fig. 4B; Param. 1 / Param. 2), suggesting that JAK2 more strongly activates STAT1 than JAK1 upon IL-6 stimulation. Building on this finding we then sought to perturb JAK2-related parameters to better understand how inhibition of JAK2 would impact downstream responses.

### Computational prediction of STAT-specific and global gene-level responses to JAK2 inhibition

To predict the impact of JAK2 perturbation on STAT phosphorylation and gene expression, we varied the individually constrained JAK2 concentration parameter (JAK2_0; Fig. S3A) and ratio-constrained IL-6-JAK2-to-STAT1 activation rate parameter (il6_jak2_med_STAT1_act; Fig. 4B). Starting from a randomly selected parameter set in our ensemble, we tested the effects of reducing either or both JAK2-associated parameter values 3-fold on IL-6-induced pSTAT dynamics (Fig. 5A). Both JAK2 parameters had a large impact on pSTAT1 amplitude (Fig. 5B, bottom row) while exhibiting more modest but also distinct effects on pSTAT3 amplitude and timing (Fig. 5B, top row).

**Figure 5.**
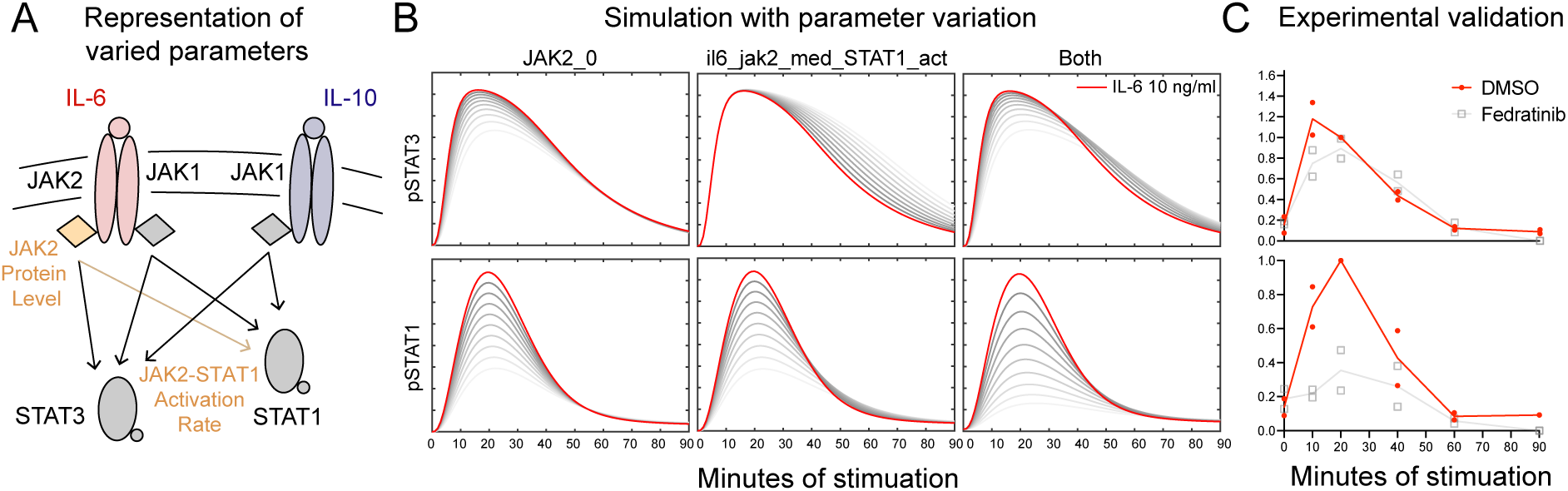
Mechanistic model prediction and experimental validation of STAT1 biased sensitivity to JAK2 inhibition. (A) Focused model schematic denoting JAK2-related model parameters in orange. (B) Impact of up to 3-fold decrease (grayscale) in JAK2 protein levels (JAK2_0), IL-6-induced JAK2 activation of STAT1 (il6_jak2_med_STAT1_act), or both parameters together on IL-6-induced (10 ng/ml) pSTAT model trajectories. The unperturbed, base parameter set is shown in red. (C) pSTAT IF data of IL-6-stimulated (10 ng/ml) BMDM, with and without 1 nM Fedratinib (JAK2-specific inhibitor) treatment (n = 2).

To validate these predictions, we stimulated BMDM with 10 ng/ml of IL-6, with and without Fedratinib, a JAK2-specific inhibitor [27], and measured pSTAT1 and pSTAT3 over time via IF (Fig. 5C). As predicted, we found that JAK2 inhibition greatly reduced the peak of STAT1 phosphorylation and had a modest impact on pSTAT3, the latter of which has been previously reported [28]. As also predicted by our mechanistic model, the JAK2 inhibitor shifted the pSTAT3 response slightly to the right (Fig. 5B-C; top row). In contrast, the literature-based model predicted that both IL-10-induced pSTAT responses would be unchanged with JAK2 parameter variation. Our experiments showed that Fedratinib reduced the IL-10-induced pSTAT1 response (Fig. S4A), suggesting the existence of undescribed signaling mechanisms involving JAK2 in the pSTAT response to IL-10.

Utilizing our full computational workflow (Fig. 1A), we trained the machine learning model with unperturbed pSTAT1 trajectories (full timecourse) to predict gene expression in response to IL-6 or IL-10 (Fig. S4B; left). We then tested our model using pSTAT1 trajectories with perturbation of JAK2 mechanistic model parameters, as discussed above (Fig. 6A). Because we trained the model to predict gene expression from pSTAT1 trajectories, the RMSE reflects the predicted magnitude of change in the expression of each gene upon JAK2 inhibition. (Fig. 6A-B). Maintaining the gene order from the hierarchical clustering of the RNAseq data (Fig. S4B; left), we predicted clusters of genes that had a median RMSE below the 10^th^ percentile would be “JAK2-independent” and those above the 90^th^ percentile would be “JAK2-dependent” (Fig 6B, Fig. S4B); the percentiles were calculated using RMSEs from all IL-6 or IL-10 gene predictions to define the 10^th^ and 90 percentiles for each cytokine, separately. We opted to evaluate groups of genes rather than individual genes that met the threshold, speculating that clusters of genes exhibiting similar dynamic behaviors may have more meaningful prediction scores.

**Figure 6.**
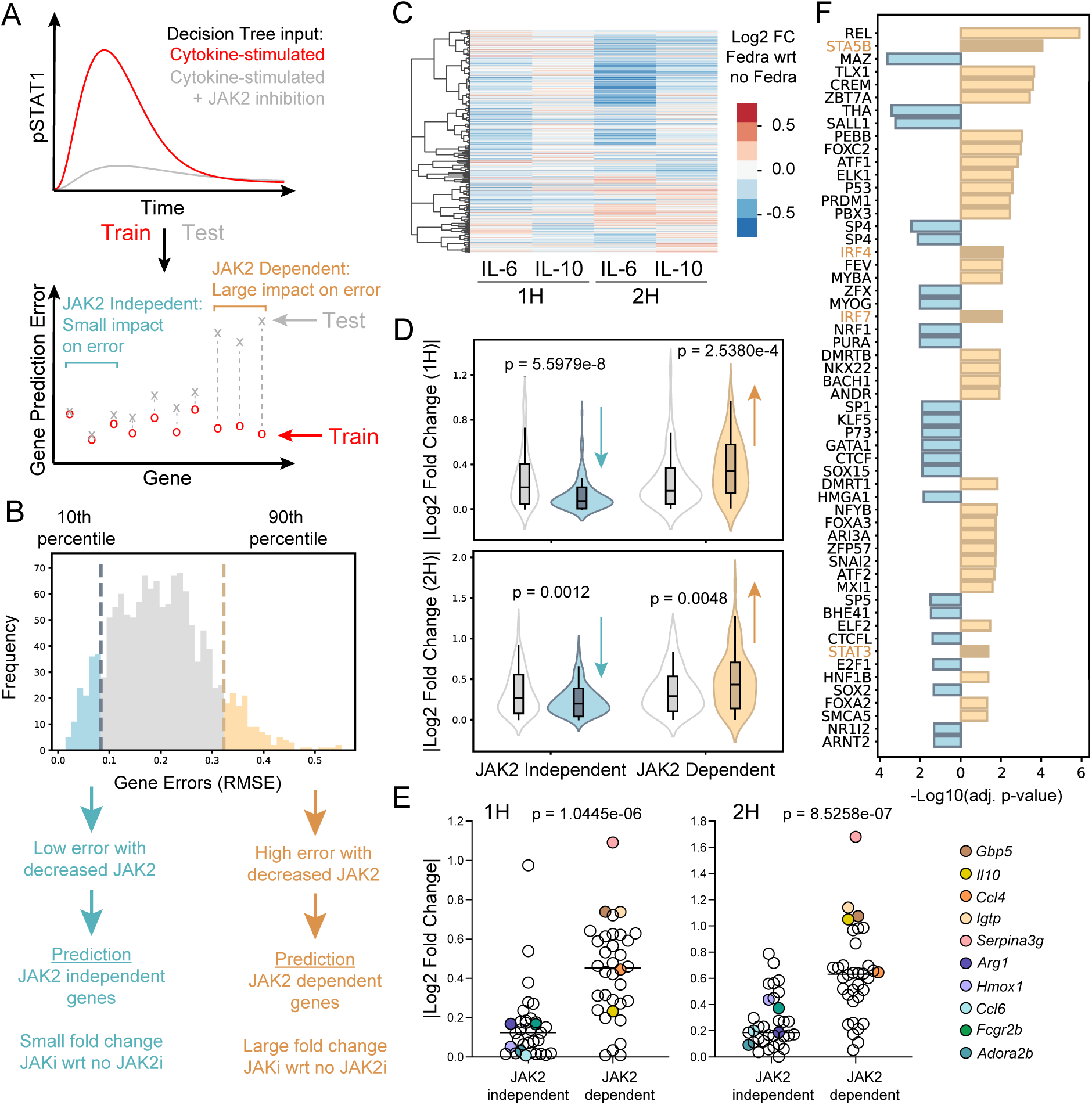
Computational prediction of global gene-level responses to JAK2 inhibition. (A) Decision Tree Regressor training and testing procedure for identification of JAK2-depedent and independent genes. (B) Model-predicted, gene expression errors (RMSE) with JAK2 parameter variation to simulate JAK2 inhibition. (C-F) BMDM were stimulated with IL-6 or IL-10, with or without 1 nM Fedratinib (JAK2-specific inhibitor) for 1h and 2h, and gene expression was quantified by RNAseq. (C) Hierarchical clustering of Log_2_ fold changes, showing cytokine-simulated (1 ng/ml) BMDM DEGs, comparing Fedratinib (1 nM) with respect to cytokine-stimulated, no Fedratinib controls; pooled samples (n = 2). (D) Absolute value of Log_2_ fold change for IL-6 + Fedratinib treated with respect to IL-6 only; predicted JAK2-dependent and predicted JAK2-independent genes were compared to randomly selected genes. (E) Absolute value of Log_2_ fold change between Immune Response genes identified in the predicted JAK2-dependent versus predicted JAK2-independent gene sets. Example genes from each group are labeled. (F) TFs significantly enriched in predicted JAK2-dependent versus predicted JAK2-independent gene sets. Highlighted motifs are STAT-related (canonical and non-canonical) transcription factors motifs (GAS and ISRE, respectively). Adjusted p-values for (D) and (E) were determined using a one-sided Welch’s t-test.

To validate our predictions, we generated RNAseq data in which we treated BMDM with 1 ng/ml of IL-6 or IL-10 at 1 and 2 hours, with and without Fedratinib (Fig. 6C). As expected, based on literature consensus that JAK2 is required for the function of IL-6 but not IL-10 [22], we saw a greater reduction in IL-6-induced gene expression, compared to IL-10-induced (Fig. 6C). For IL-6-induced genes, we then compared our computationally predicted JAK2-dependent and JAK2-independent genes, plotting their absolute value Log_2_ fold changes in the validation Fedratinib RNAseq data (Fig. 6D). The computationally predicted JAK2-independent genes had significantly lower fold changes, while the predicted JAK2-dependent genes had significantly higher fold changes, both compared to a gene random set of the same size (Fig. 6D). For IL-10-stimulated genes, our model did not accurately predict fold change in response to Fedratinib (Fig. S4C), consistent with undescribed and unmodeled JAK2-dependent IL-10 mechanisms (Fig. S4A). For IL-6 induced genes, we ran a gene set enrichment analysis (GSEA; [29, 30]) on the predicted JAK2-dependent and JAK2-independent genes; the Immune Response biological process was significantly enriched in both gene sets. When we plotted the Log_2_ fold changes of Immune Response genes, we found the fold changes of genes predicted as JAK2-dependent were significantly increased compared to those predicted as JAK2-independent (Fig. 6E) further validating our JAK2 sensitivity predictions.

Our results suggest there exist gene-specific mechanisms for decoding STAT dynamics that lead to differential gene sensitivity to JAK2 inhibition. To investigate this possibility, we scanned promoter regions of either our predicted JAK2-dependent or JAK2-independent genes to identify TF motifs that were significantly enriched in one gene set over the other. We found 33 TF motifs that were significantly enriched in the JAK2-dependent gene set, while 23 were enriched in the JAK2-independent gene set (Fig. 6F). We noted that canonical STAT binding motifs (GAS; i.e., STAT5B and STAT3; Fig. S4D) and non-canonical STAT binding motifs (ISRE; i.e., IRF4 and IRF7) were enriched in the JAK2-dependent genes. Consistent with direct JAK2 regulation of STATs, this may suggest disparate STAT dependence between the two gene sets. Additionally, JAK2 may regulate other TFs that support the decoding of STAT dynamics and the diversity in gene induction. Overall, our results demonstrated that the integrated workflow can make meaningful predictions of the effects of kinase inhibition on TF dynamics and gene expression.

## Discussion

Linking signaling events to the expression of specific gene sets and associated functional outcomes is critical for rational therapeutic targeting of signaling regulators. Here, we proposed a mechanistic-to-machine learning model of cytokine-induced JAK-STAT signaling in macrophages to demonstrate the ability of systems modeling approaches to link quantitative perturbation of upstream signaling network components to varied effects on global downstream gene expression. Our integrated model allowed us to link previously appreciated cytokine-specific STAT3 dynamics to genes that decode sustained STAT3 activity and predicted sensitivity to JAK2 inhibition, both at the level of STAT activity and cytokine-induced gene sets. We demonstrated the value of such a modeling framework for making predictions at multiple levels of a signaling-to-transcription network to our understanding of how cellular response diversity arises.

Prior work in the temporal coding theory has focused on various portions of the context-specific, signaling-to-gene expression process. Some studies have focused on detailed modeling of upstream signaling events, associating these events with stimulus-specific gene expression profiles on a relatively small scale. For example, studies have linked key TF dynamic features to specific transcriptional patterns in a small number of genes, primarily focusing on the upstream signaling events and mechanisms [11, 13, 31]. These studies provide an in-depth understanding of mechanisms shaping the induced dynamics but provide only a glimpse of the downstream outcomes. Other studies have worked to connect stimulus-specificity to more global gene expression signatures, finding that TF dynamics can encode specific information about the activating stimulus, and then influence gene expression profiles downstream [10]. While this allows us to understand that altered TF dynamics can be misinterpreted, resulting in dysregulation at the gene level, it still does not allow us to connect specific signaling events to sets of genes. A major benefit of our integrated modeling approach is the ability to address context-specificity at multiple levels within the signaling-to-gene expression process; our approach allows us to perturb signaling components and predict outcomes globally at the level of gene expression.

Our workflow depends on interpreting the variation of signaling components in the mechanistic model. As mechanistic signaling models grow in complexity, the large number of unknown parameters, relative to the amount of data available for fitting, makes models difficult to interpret. Estimating parameters of such large models leads to issues with practical identifiability, or the assumption that consistent parameter values can be estimated from experimental data [32]. Given that acquiring sufficient data to make model parameters identifiable is often not feasible, resulting in situations in which parameter distributions are not well constrained, there is a significant need to develop parameter analysis approaches to extract quantitative biological insights from model results. To gain some insights from our under-constrained model, we conducted a parameter correlation analysis post-fitting, which uncovered some constrained signaling relationships, such as the relative JAK1 and JAK2 activation of the STATs. Although the individual parameters were not sufficiently constrained by the experimental data, we were still able to make predictions about the relative JAK activity on gene expression. The parameter analysis here utilized simple correlations to identify relationships between various model components, but more complex analysis of latent factors or the utilization of non-linear approaches may provide additional insight into larger groups or modules of parameters whose coordinated regulation is associated with downstream response specificity. This parameter analysis is just the first step in the development of other methods to identify signaling modules that can uncover complex parameter relationships in under-constrained signaling models.

TYK2, JAK1 and JAK2 have all been established as activating kinases for the IL-6 receptor complex [22]. In contrast IL-10 requires JAK1 for its anti-inflammatory function and it has been reported that IL-10 induces the phosphorylation of TYK2 and JAK1, but not JAK2 or JAK3 [22–24, 33]. Consistent with these known mechanisms, our literature-informed signaling model accurately predicted the effects of JAK2 inhibition on IL-6-induced signaling and gene expression. Given our assumption that JAK2 is not directly involved in the IL-10-induced response, it was not surprising that our model predicted no impact of JAK2 inhibition on the IL-10-induced dynamic response. In contrast, our experimental data showed an impact of JAK2 inhibition on IL-10-induced STAT phosphorylation and gene expression profiles. Though these effects were modest compared to those of IL-6, the data suggests a role of undescribed signaling mechanisms in shaping IL-10-induced gene expression. Numerous non-mutually exclusive mechanisms could contribute to non-predicted features in our data, such as IL-10-induced genes whose expression increased with JAK2 inhibition; possibilities include competition between JAKs at the level of cytokine receptors or variation in the composition of STAT dimers, which were not explicitly modeled. The literature has noted that IL-6-induced STAT1:STAT1, STAT1:STAT3, and STAT3:STAT3 dimers form at different rates and exhibit unique nuclear translocation efficiencies [6], but mechanisms surrounding the STAT dimers and their associated functions are unclear. Our data suggest that JAK2 is required for the modest STAT1 activation induced by IL-10, compared to IL-6, and are consistent with the possibility that the absence of pSTAT1 and thus, STAT1:STAT3 heterodimers, leads to enhanced IL-10-induced STAT3 homodimer formation and STAT3 dependent gene expression with JAK2 inhibition. Future iterations of integrated JAK-STAT models can aim to incorporate the different mechanisms above to generate testable hypotheses and enhance our ability to predict IL-10-induced gene expression.

Integrated systems modeling approaches such as ours have shown the inherent value of multi-level modeling for learning about the mechanisms contributing to diversity in signaling and downstream gene expression. Although we successfully demonstrated this in the IL-6-induced JAK-STAT response, the incorporation of single-cell dynamics and gene expression data has the potential to significantly improve the power of such methodology. While our single-cell phosphorylation data suggest that IL-6 and IL-10 do not induce bimodal pSTAT responses, cell-to-cell variation in the magnitude of STAT phosphorylation at each timepoint likely contributes to variation in gene expression at the single-cell level. Multiple studies support the idea that cell-to-cell variation in gene expression guides appropriate cellular population responses to complex, and even contrasting, environmental stimuli [34–36]. Furthermore, in the NF-kB system, varying patterns of single-cell TF dynamics have been associated with different single-cell gene expression profiles downstream [18, 36, 37]. Tracking single-cell TF dynamics and gene expression together over time would likely improve the identifiability of our mechanistic model parameters and support more accurate machine learning predictions, regarding the impact of signaling variation on downstream gene expression. We utilized a simple and highly interpretable machine learning model to make gene predictions, given that our stimulus-specific simulated data followed relatively consistent activation patterns. The addition of single-cell readouts and the resulting heterogeneity would enable us to use more complex machine learning methods, such as recurrent neural networks, which are time varying in nature [38] or even causal modeling methods [39], which would improve our predictive capabilities and give us more biological insights. Beyond these technical advantages, connecting single-cell signaling variation to gene expression readouts would allow us to quantify response heterogeneity and underlying mechanisms, in particular, those that may arise from treatment with various inhibitors [40].

Cytokine-induced gene expression is largely attributed to the JAK-STAT signaling system. Multiple studies have shown STAT3 to be necessary for appropriate cytokine-induced functional responses [24, 41, 42], although it is well-appreciated that other TFs support cytokine-specific function. STATs canonically bind to gamma-activated sequence (GAS) DNA motifs to induce gene expression [2], but the interaction of STATs with various IRFs can shift the DNA binding profile towards the IFN-stimulated response element (ISRE) binding, resulting in altered gene induction [7]. NF-kB and STATs notably interact both directly and indirectly to alter gene expression outcomes upon cytokine sensing, further contributing the context-specific diversity in STAT responses [8, 43]. Even STAT1 and STAT3 specifically, interact with each other to enhance the diversity of downstream gene expression [17]. Taken together, the JAK-STAT pathway is central to cytokine-specific gene expression, although interactions with other signaling pathways and TFs undoubtedly support STAT specificity and diversity. Our motif analysis is consistent with involvement numerous non-STAT TFs to support sensitivity of gene sets to cytokine-specific STAT dynamics and JAK inhibition. Efforts to validate factors regulating these context-specific gene sets would greatly enhance our ability to predict and perturb cytokine regulated cellular function. For example, ChIP-seq data would yield insight into the gene specific binding of candidate STAT cooperating TFs. Additionally, given the likely contribution of chromatin accessibility to the responsiveness of STAT target genes to upstream signaling dynamics, ATAC-seq would enhance our mechanistic understanding of signal decoding mechanisms. Integrating such datasets with our signaling dynamics would allow us to improve upon our predictive gene expression model by constraining graph structures of more powerful machine learning models with biological data [44], allowing us to infer more complex regulatory mechanisms supporting cytokine-specific gene induction.

To our knowledge, we are the first to systematically identify gene sets in global transcriptomic data predicted by dynamic signaling features and kinase perturbation. Our mechanistic-to-machine learning model of JAK-STAT signaling from initiation to gene expression serves as a broadly applicable template for interrogating diverse dynamic signaling systems. While we have taken a forward approach, by predicting TF dynamics, and subsequently, cytokine-induced gene sets, future efforts can take a reverse approach, beginning with cytokine-induced gene sets of interest and inferring signaling mechanisms supporting their induction. In summary, this modeling workflow improves our mechanistic understanding of cytokine-specificity and serves as a fundamental proof of concept and framework to guide more comprehensive modeling studies of complex cell signaling-to-gene expression systems.

## Methods

### Experiments and quantification

#### Mice, Macrophage culture, and cytokine stimulation

Female C57BL/6J mice (8-12 weeks old) were purchased from Jackson Laboratory and maintained in specific-pathogen-free conditions. All procedures were approved by the Institutional Animal Care Committee of the University of Pittsburgh (IACUC). To generate bone marrow derived macrophages (BMDMs), mouse bone marrow was cultured for 6 days in Dulbecco’s modified Eagle’s medium (DMEM) with 10% fetal bovine serum (FBS), penicillin (100 U/ml), streptomycin (100 U/ml), 2 mM l-glutamine, and 20 mM Hepes supplemented with recombinant mouse M-CSF (60 ng/ml; R&D Systems). BMDMs were scraped with cold PBS and plated in either a 48 or 96 well plate for qPCR and IF, respectively, and rested overnight at 37°C. BMDMs were stimulated on day 7 with the appropriate doses of IL-6 (R&D Systems) and IL-10 (R&D Systems) at the noted time points. In some experiments, cells were treated with 1 nM of Fedratinib 20 minutes prior to cytokine stimulation.

#### Immunofluorescence imaging and analysis

Stimulated cells were fixed for 10 minutes with 4% paraformaldehyde (PFA) at room temperature and permeabilized with 100% methanol for an additional 10 minutes at -20°C. Cells were then blocked in PBS with 5% goat serum and 0.3% Triton X-100 for one hour at room temperature. Cells were stained with primary antibodies diluted in PBS with 1% bovine serum albumin (BSA) and 0.3% Triton X-100 at 4°C overnight. The following primary antibodies were used: anti-CD11b (BioRad MCA711, 1:400), anti-pSTAT3 Y705 (Cell Signaling Technology 4113, 1:200), and anti-pSTAT1 Y701 (Cell Signaling Technology 9167, 1:400). Cells were then washed and stained for 1 hour at room temperature with the following secondary antibodies: anti-hamster Cy3 (Jackson ImmunoResearch 127165160, 1:500), anti-rabbit AlexFluor 488 (Invitrogen A11008, 1:500), and anti-mouse AlexFluor 647 (Invitrogen A21235, 1:500). Cells were lastly stained with Hoechst 3342 (Thermo Fisher Scientific). The cells were imaged and analyzed with the CellInsight CX5 High Content Screening Platform and HCS Studio (Thermo Fisher Scientific).

Mean fluorescent intensity (MFI) of phospho-STATs were measured over whole cells, defined as objects using surface CD11b staining. Experiments were scaled by shifting starting MFI values (time = 0) to zero and setting any resulting negative values to 0. To support pooling across experiments, all data points within a given experiment were normalized to the IL-6 10 ng/ml 20 minute time point MFI value; pSTAT1 and pSTAT3 were normalized independently. To get the means of each stimulation condition that would later be used for model fitting, Python’s SciPy [45] interpolation function was used to fill in missing time points in select experiments. pSTAT3 and pSTAT1 average timecourses for each stimulation condition were stacked column-wise (pSTAT3 first, then pSTAT1), and conditions that produced similar dynamics behaviors were clustered together using K-means clustering from the scikit-learn package [46] with default settings.

#### RNAseq analysis

Cells were lysed with TRIzol Reagent (Ambion) and RNA was isolated with the Direct-zol RNA MicroPrep Kit (Zymo Research). The Illumina Stranded mRNA Prep protocol was used with an input of 90 ng/ml per sample to prepare double stranded cDNA. Final libraries were sequenced on the NextSeq2000 (Illumina) with a loading concentration of 750 pM to a depth of 20 million, 100 base pair, paired-end reads. The Mus musculus reference genome (mm10) was downloaded from the UCSC genome browser [47, 48], and Rsubread [49] was used to align and obtain raw read counts with that reference. RNAseq files were submitted to Gene Expression Omnibus (GSE231345). DESeq2 [50] was used for read normalization and differential gene expression calculations. P-values were corrected using the Benjamini-Hochberg method for multiple hypothesis testing. Genes with expression averages above a 10 read threshold and those that were differentially expressed in any condition (adjusted p-value ≤ 0.05) were included in further analysis. For the RNAseq data shown in Figure 3, including 1 ng/ml and 10 ng/ml of IL-6, IL-10, and combined IL-6 and IL-10-stimulated BMDM at 1, 2, and 4 hours, genes with a fold change of more that 1.5 were included in subsequent analysis and were z-score normalized. For the Fedratinib-treated validation RNAseq dataset shown in Figure 6, we removed genes that had a high variance across replicates and calculated the Log_2_ fold change for the remaining genes by comparing the cytokine-stimulated inhibitor-treated samples with cytokine-stimulated control samples. RNAseq heatmaps were clustered using hierarchical clustering using correlation as the distance metric and complete linkage.

## Mechanistic modeling

### Model architecture and construction

The JAK-STAT mechanistic model was adapted from a previous IL-6 and IL-10-induced pSTAT3 model in dendritic cells [13], with integration of the separate IL-6 and IL-10 models. We also added STAT1 phosphorylation, an additional SOCS protein, and two generic phosphatase negative regulators. Our model was coded as a rule-based model using the BioNetGen language (BNGL) [21]. The model was simulated using the ordinary differential equations (ODEs) from 0 to 90 minutes, with one minute time step intervals. The BNGL and accompanying files to simulate the model in MATLAB can be found at https://github.com/ncheemalavagu/STAT_models.

### Parameter estimation (model fitting)

We estimated unknown model parameters from experimental pSTAT1 and pSTAT3 IF data using a MATLAB implementation of parallel tempering called pTempEst, a Bayesian parameter estimation approach with an enhanced sampling method [25]. Parallel tempering more efficiently samples high dimensional parameter space by running multiple Markov chains simultaneously but at different temperatures. Higher temperature values scale the energy landscape to allow for the chain to escape local minima more easily and therefore, increase the efficiency of sampling. The normalized responses of IL-6, IL-10, and the combination of the two at 1 ng/ml and 10 ng/ml were averaged for the model fitting, and the standard deviation was set to 15% percent of the mean as an estimate of biological variation observed in our data. We assumed all our unknown model parameters to be log-normally distributed for the parameter estimation setup. Since our data used for fitting was normalized to the IL-6 10 ng/ml 20 minute time point for STAT1 and STAT3 separately, model fits were also scaled accordingly prior to calculating the log-likelihood to evaluate the current model fit. We used three Markov chains for pTempEst parameter sampling, and the rest of the settings were kept to the default. The initial parameter chains were randomly initialized and approximately 110,000 chains swaps were needed for the chains to stabilize at a low energy value. After that point, parameter sets that allowed our model simulations to fit the experimental data without oscillatory behaviors were selected for further analysis.

### Parameter analysis and modulation

To evaluate the constraint of our estimated model parameter values, we bootstrapped 50 samples across 5000 runs from each of our parameter distributions and used the Shapiro-Wilk test on the log-transformed values to determine whether the distribution could be a normal distribution as expected for a well-constrained parameter. If half or more of the bootstrapped runs had a false discovery rate greater than 0.05 (the alternative hypothesis suggests normality) and if the parameter values were within a 2 log range, the parameter was considered log-normally distributed and well-constrained by the experimental data. With the remaining, under-constrained parameters, we calculated the correlation coefficients between them and used hierarchical clustering (Euclidean distance metric, complete linkage) to group together parameters that compensated for each other. To model JAK2 inhibition in the mechanistic model, we first selected the parameter set that fit our data the best after parameter estimation (lowest energy value). After confirming that the JAK2 concentration parameter (JAK2_0) and the IL-6-JAK-to-STAT1 activation rate ratio (il6_jak1_med_STAT1_act:il6_jak2_med_STAT1_act) were in the middle of the constrained distributions, we reduced the value of the JAK2-associated parameters individually and together, otherwise fixing the parameter values, for up to a 3-fold reduction and simulated pSTAT1 and pSTAT3 trajectories to make STAT dynamic predictions to JAK2 inhibition.

## Machine learning prediction and validation

### Timeframe identification

We constructed at 7-state Gaussian-emission Hidden Markov Model (HMM) using the hmmlearn API to identify dynamic timeframes from mechanistic model simulated pSTAT3 trajectories. Using the parameter sets from parameter estimation, we split the data into approximately 16,000 simulated trajectories for training and 1500 simulated trajectories for testing, both of which consisted of equal numbers of IL-6, IL-10, and combo-stimulated (low and high dose) responses. Random initialization was used, and the initial state was always in state 1 because all pSTAT3 trajectories begin at 0. The transition limitations are noted in Fig. S2A. We sampled 250 bootstrapped samples across 10,000 runs to estimate the transition matrix and emissions (means and covariances) using the Baum-Welch algorithm. The final HMM transition matrix and emission values were averaged across the bootstrapped runs and used for testing. The testing pSTAT3 trajectories were then used to identify the state changes for each timecourse using the Viterbi algorithm. The procedure gave us a sequence of states (1-7) for every time point (0 - 90 minutes) in each timecourse (n = 1500). To identify timeframes of the pSTAT3 dynamics response (Fig. S2B), we noted the time at which a dynamic state began and the time at which a state switch occurred for each timecourse. Then, we averaged the starting and ending points across the 1500 trajectories to identify timeframes. Overlapping portions of two timeframes were considered their own separate frame. Gene expression was predicted with decision tree regressors (details below) using the identified timeframes, and timeframes with highly correlated gene predictions were grouped to define our early and late timeframes.

### Decision tree regressor gene prediction

We trained and tested decision tree regressors with the scikit-learn implementation [46] and default settings using the same data split as the HMM above. The input data to the decision tree consisted of the pSTAT trajectories (pSTAT3 for Fig. 3 analysis and pSTAT1 for Fig. 6 analysis). The corresponding labels were flattened, gene expression data for the corresponding stimulation condition (i.e., IL-6 1 ng/ml trajectory with IL-6 1 ng/ml gene expression). The gene expression values for decision tree predictions were mean centered to the control sample. For the early versus late pSTAT3 gene predictions (Fig. 3), the decision tree regressor model was trained and tested using the appropriate portion of the trajectory, and for the pSTAT1 JAK2 inhibitor gene predictions (Fig. 6), the model was trained using the full, unperturbed pSTAT1 trajectory, and tested with the full, JAK2-inhibited pSTAT1 trajectory.

### Early versus late timeframe prediction of cytokine-specific clusters

RNAseq gene clusters were identified as cytokine-specific if they were induced (adj. p-value <= 0.05) by a single cytokine over the three timepoints. This comparison was done for the high and low doses independently. If at any time point or dose, the opposing cytokine also induced a particular gene, that gene was not considered cytokine specific. Using hierarchical clustering (correlation distance metric, complete linkage), we selected the number of total gene clusters that maximized the number of cytokine-specific clusters. A cluster was considered cytokine-specific if at least 1/3 of the genes were cytokine-specific based on the above criteria and only cytokine-specific clusters of at least 10 genes (sufficiently large) were considered for further analysis.

### JAK2 inhibitor prediction experimental validation

To validate our JAK2-dependent and JAK2-independent gene predictions using RNAseq in the context of Fedratinib treatment, we averaged RMSEs of cytokine-induced gene predictions across the 1 and 2 hour time points (the timepoints for which we had Fedratinib RNAseq validation data). Genes were considered for validation only if they were upregulated by cytokine in both the Fedratinib and original single cytokine RNAseq datasets. For evaluating our predictions of JAK2-dependent and independent genes, we used one-sided Welch’s t-tests to determine whether JAK2-independent genes had significantly lower fold changes relative to the random set and if JAK2-dependent genes had significantly higher fold changes relative to the random set. To assess predictions more directly for immune-relevant genes, gene set enrichment analysis of mouse Gene Ontology terms was conducted on IL-6-stimulated, predicted JAK2-dependent and predicted JAK2-independent gene sets. The GOBP_Immune_Response term was enriched in both gene sets. For GOBP_Immune_Response genes from the predicted JAK2-dependent and predicted JAK2-independent gene sets, we compared their Log_2_ fold changes with Fedratinib-treated IL-6 samples, with respect to IL-6-induced without inhibitor treatment. Again, a one-sided Welch’s t-test was used to test whether the JAK2-independent GOBP_Immune_Response genes had significantly lower fold changes than the JAK2-dependent GOBP_Immune_Response genes.

## Transcription factor motif analysis

To identify transcription factor motifs enriched in the JAK2-dependent versus independent gene groups, promoter binding regions were obtained from the UCSC genome browser mm10 build and defined as 500 base pairs downstream to 100 base pairs upstream of the gene transcriptional start sites. PWMscan [51] was used to search the defined promoter regions for each group for given binding motifs. We scanned for all mouse transcription factor motifs from the HOCOMOCO database [52]. Once we had the number of hits for each motif in the defined promoter regions, we removed motifs in overlapping regions and used a Fisher’s Exact Test to identify motifs that were enriched in one gene set over the other. The number of promoter regions scanned that did not contain a particular motif was used as the background. To correct for testing multiple motifs, we used the Benjamini-Hochberg multiple hypothesis correction to adjust our p-values. An adjusted p-value of less than or equal to 0.05 was considered statistically significant.

## Acknowledgments

Thank you to members of the Gottschalk and Faeder labs, and colleagues from the Center for Systems Immunology for discussion and critical feedback. This work was supported the National Institute of General Medical Sciences 1R35GM146896. Neha Cheemalavagu was supported by 5T32-AI1089443 and James R. Faeder was supported by P41GM10371.

## Figure legends

## List of Supplemental files

**Table S1. Table S1. Rule-based model parameter names and descriptions.**

**Supplemental File 1. Genes in late pSTAT3 predicted clusters noted by asterisks in Fig. 3C.**

**Figure S1.**
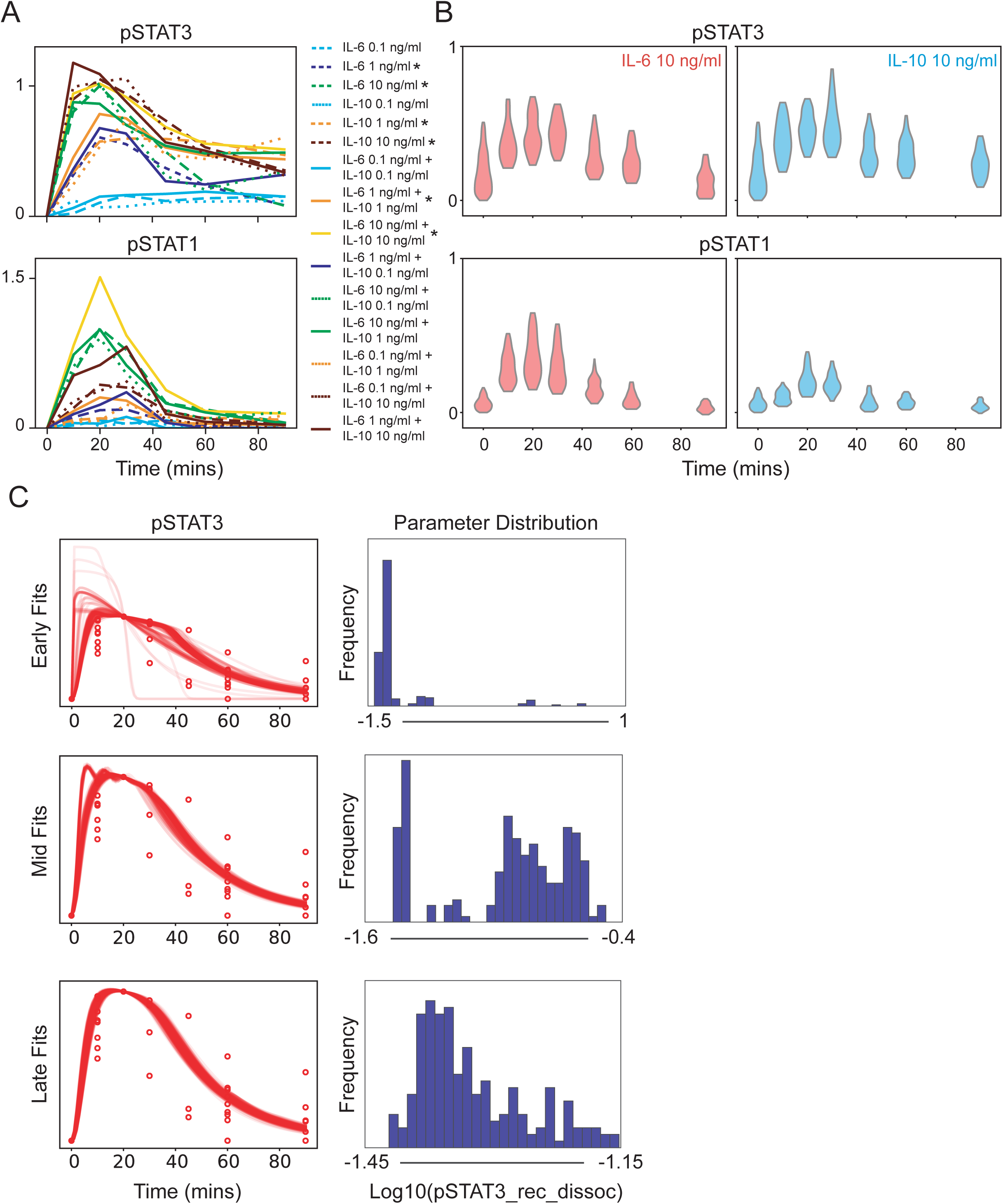
Detailed analysis of pSTAT dynamics and model fitting steps. (A) Cytokine-induced pSTAT dynamic curves colored according to K-means clustering results. Asterisks note conditions selected for RNAseq and all further analysis. (B) Violin plots of a representative experiment showing single-cell distributions of IL-6 or IL-10-induced (10 ng/ml) pSTAT1 and pSTAT3. (C) Parameter estimation results at early, middle (mid), and late stages of fitting process (left column) with corresponding Log_10_ of selected model parameter (right column). Detailed description of parameter estimation process in Methods.

**Figure S2.**
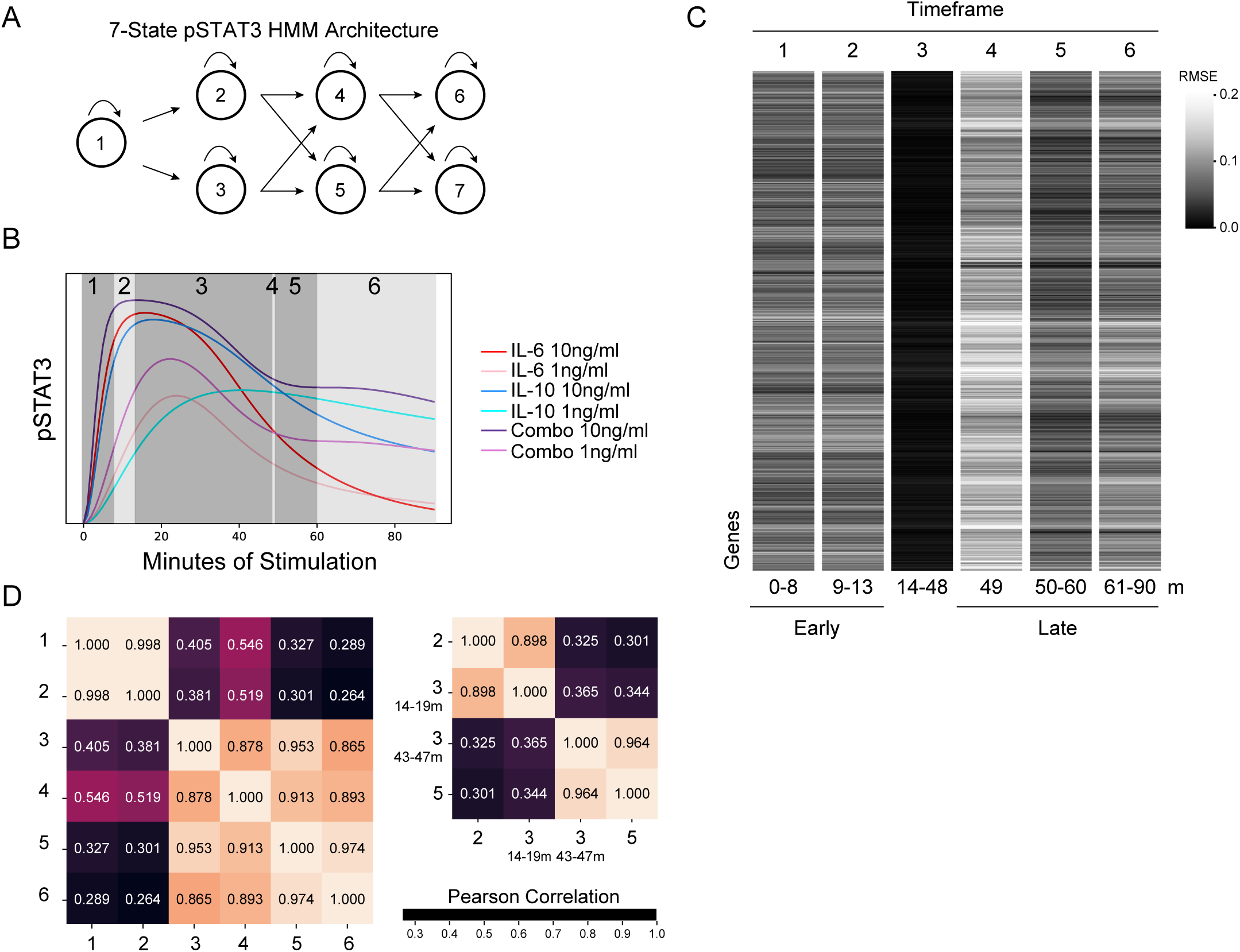
Early versus late STAT3 dynamic timeframes identified using a Hidden Markov Model (HMM). (A) 7-state HMM architecture used to identify dynamic STAT3 phosphorylation states (hidden states). Gaussian emission HMM was trained with the Baum-Welch algorithm on 10,000 bootstrapped runs with 250 model-simulated, cytokine-induced pSTAT3 samples each, and the transition matrix, means, and covariances of the states were averaged across runs for the final HMM. The HMM hidden states were decoded in test pSTAT3 trajectories using the Viterbi algorithm. (B) 6 pSTAT3 timeframes defined by HMM. Boundaries of timeframes were identified by selecting the time points at which each trajectory underwent a transition to a new state (from Viterbi decoding), and those time points were averaged across all cytokine-induced trajectories to be used as timeframe boundaries. The overlapping portions of timeframes were considered their own timeframe. (C) Gene expression prediction errors (RMSE) averaged across time points per timeframe using the prediction pipeline in Fig. 3A. (D) Correlation matrices of timeframe gene predictions, demonstrate that timeframes 1 and 2 exhibit similar prediction behaviors, while timeframes 3 through 6 exhibit similar prediction behaviors (left matrix). Correlations of the early and late portion of timeframe 3 suggest that it contains dynamic information related to the early and late responses (right matrix). Timeframes 1 and 2 correspond to “early” in Figure 3, while timeframes 5 and 6 correspond to “late”.

**Figure S3.**
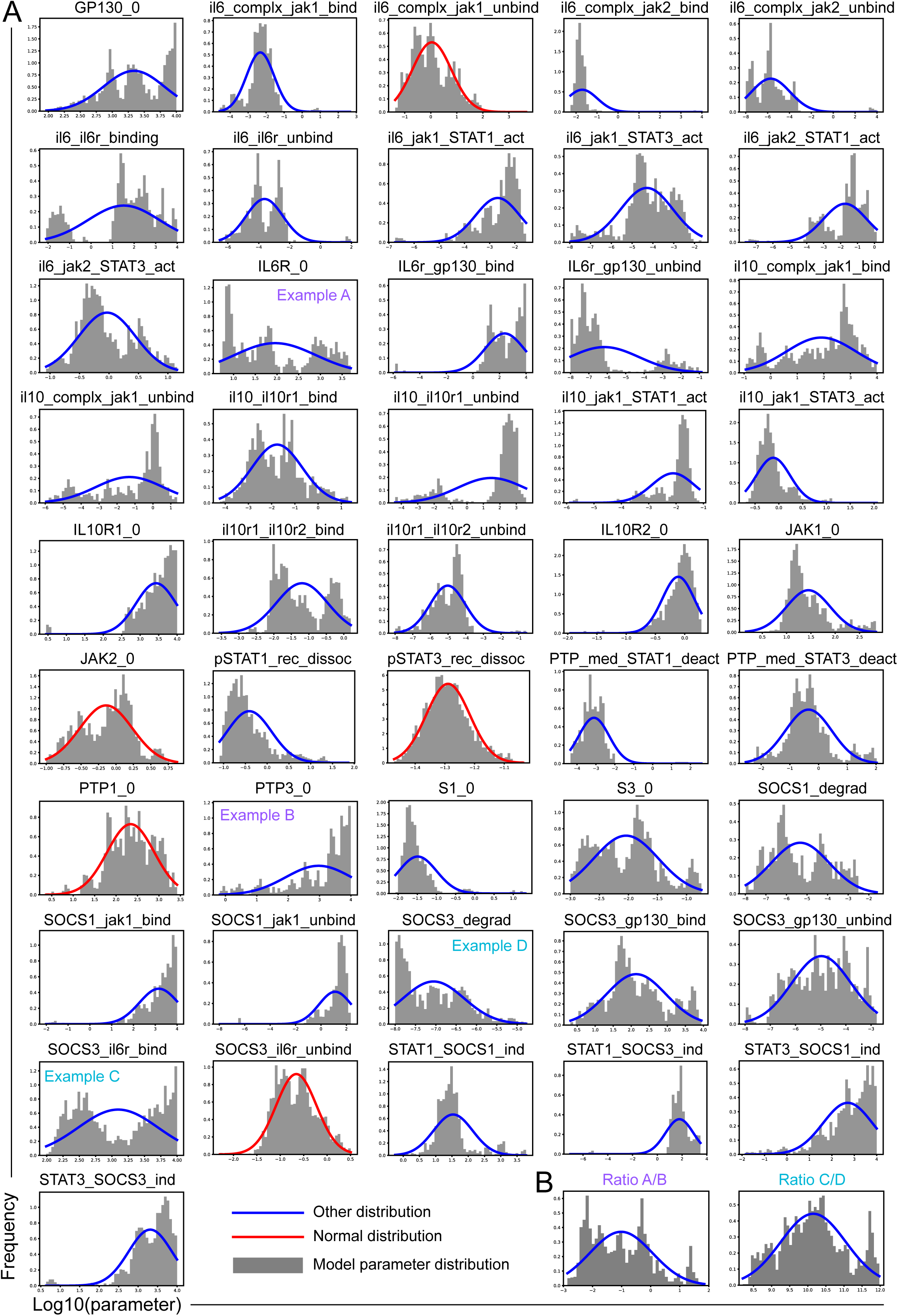
Rule-based model parameters are under constrained by experimental data. (A) Distributions of Log_10_ model parameter values. Colors represent normality testing results from the Shapiro-Wilk test, with red curves showing distributions deemed log normally distributed in more than half of bootstrapped runs. (B) Parameter ratio distributions of randomly selected unrelated model parameter pairs, for comparison to correlated parameter ratios in Figure 4B.

**Figure S4.**
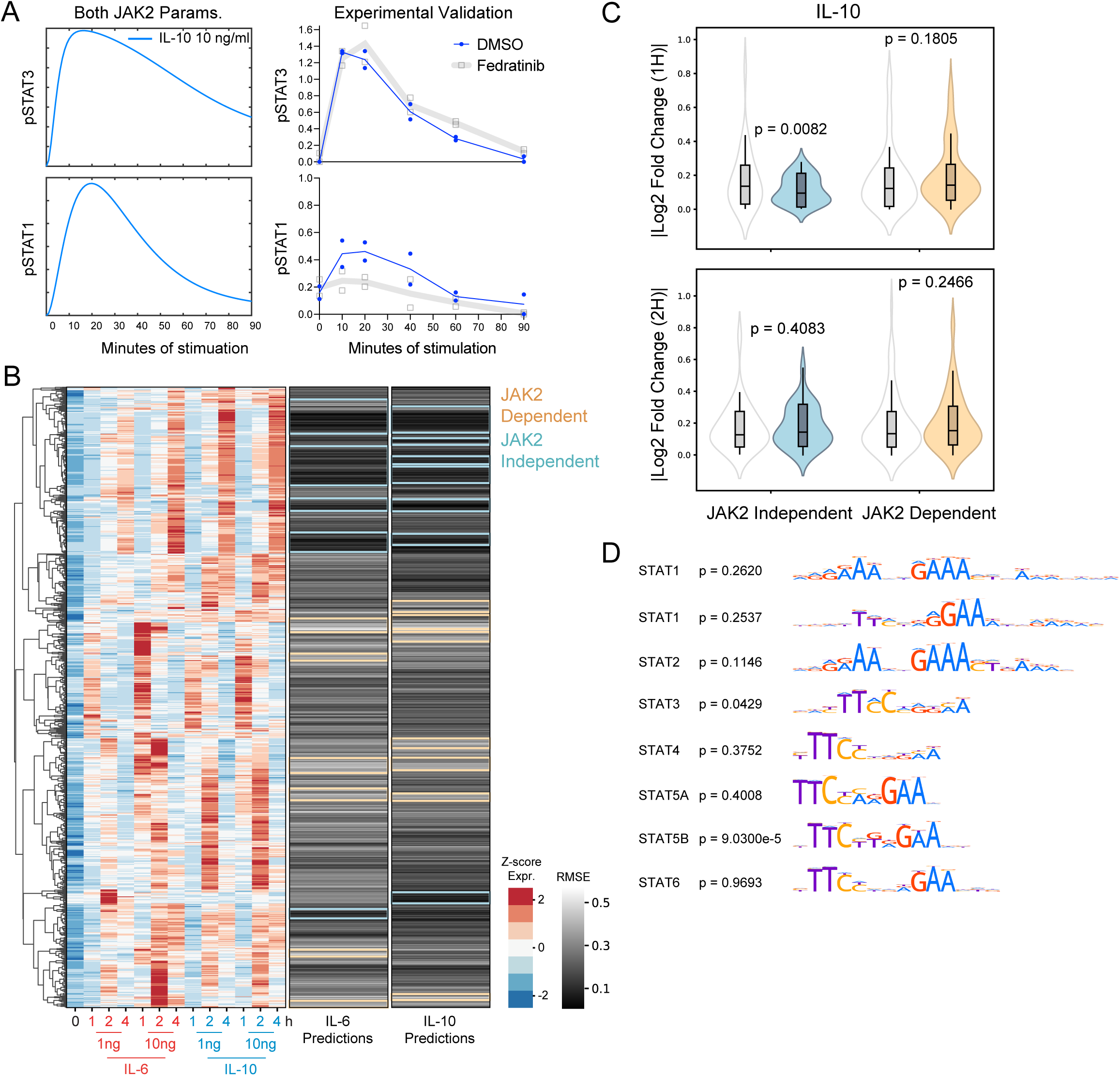
Model dynamic and gene expression predictions of IL-10-induced JAK2 inhibition are limited. (A) Impact of up to 3-fold decrease (grayscale) in both JAK2 protein levels and IL-6-induced JAK2 activation of STAT1 on IL-10-induced (10 ng/ml) pSTAT model trajectories (baseline in blue). The grayscale curves are obscured by the original blue trajectory because JAK2 does not impact the simulated IL-10-induced response. Data on the right show pSTAT IF data of IL-10-stimulated (10 ng/ml) BMDM with and without 1 nM Fedratinib treatment (n = 2). (B) Hierarchical clustering of upregulated, z-score normalized DEGs across pooled, single cytokine samples (n = 2), average gene expression prediction errors across the 1 and 2 hour, for IL-6-induced and IL-10-induced pSTAT1 gene predictions. Groups of predicted JAK2-dependent and predicted JAK2-independent genes that meet the 90th and 10th gene expression percentile cutoffs, respectively, noted by the colored boxes (see Methods for cluster identification). (C) Absolute value of Log_2_ fold change for IL-10 + Fedratinib treated with respected IL-10 only; predicted JAK2-dependent and predicted JAK2-independent genes were compared to randomly selected genes. P-values were determined using one-sided Welch’s t-test. D) Adjusted p-values and motif logos for enrichment of canonical STAT motifs in predicted JAK2-dependent versus predicted JAK2-independent genes.

