## Supplemental Table 1 for "Predicting gene level sensitivity to JAK-STAT signaling perturbation using a mechanistic-to-machine learning framework"

**Table 1. Unknown rule-based model parameter names and descriptions.**

| 1 | il6_il6r_binding | IL-6 binding to IL-6R |
| --- | --- | --- |
| 2 | il6_il6r_unbinding | IL-6 unbinding from IL-6R |
| 3 | il6r_gp130_binding | Bound IL-6/IL-6R binding to GP-130 |
| 4 | il6r_gp130_unbinding | Bound IL-6/IL-6R unbinding from GP-130 |
| 5 | il6_complex_jak1_binding | JAK1 binding to IL-6R (in IL-6 complex) |
| 6 | il6_complex_jak1_unbinding | JAK1 unbinding from IL-6R (in IL-6 complex) |
| 7 | il6_complex_jak2_binding | JAK2 binding to GP-130 (in IL-6 complex) |
| 8 | il6_complex_jak2_unbinding | JAK2 unbinding from GP-130 (in IL-6 complex) |
| 9 | SOCS3_il6r_binding | SOCS3 binding to IL-6R (in IL-6 complex) |
| 10 | SOCS3_il6r_unbinding | SOCS3 unbinding from IL-6R (in IL-6 complex) |
| 11 | SOCS3_gp130_binding | SOCS3 binding to GP-130 (in IL-6 complex) |
| 12 | SOCS3_gp130_unbinding | SOCS3 unbinding from GP-130 (in IL-6 complex) |
| 13 | il6_jak1_med_STAT3_act | IL-6 bound JAK1 phosphorylating STAT3 |
| 14 | il6_jak1_med_STAT1_act | IL-6 bound JAK1 phosphorylating STAT1 |
| 15 | il6_jak2_med_STAT3_act | IL-6 bound JAK2 phosphorylating STAT3 |
| 16 | il6_jak2_med_STAT1_act | IL-6 bound JAK2 phosphorylating STAT1 |
| 17 | il10_il10r1_binding | IL-10 binding to IL-10R1 |
| 18 | il10_il10r1_unbinding | IL-10 unbinding from IL-10R1 |
| 19 | il10r1_il10r2_binding | Bound IL-10/IL-10R1 binding to IL-10R2 |
| 20 | il10r1_il10r2_unbinding | Bound IL-10/IL-10R1 unbinding from IL-10R2 |
| 21 | il10_complex_jak1_binding | JAK1 binding to IL-10 receptor complex |
| 22 | il10_complex_jak1_unbinding | JAK1 unbinding from IL-10 receptor complex |
| 23 | il10_jak1_med_STAT3_act | IL-10 bound JAK1 phosphorylating STAT3 |
| 24 | il10_jak1_med_STAT1_act | IL-10 bound JAK1 phosphorylating STAT1 |
| 25 | SOCS1_jak1_binding | SOCS1 binding to JAK1 (in IL-10 complex) |
| 26 | SOCS1_jak1_unbinding | SOCS1 unbinding from JAK1 (in IL-10 complex) |
| 27 | pSTAT3_rec_dissoc | Phosphorylated STAT3 dissociating from receptor complex |
| 28 | pSTAT1_rec_dissoc | Phosphorylated STAT1 dissociating from receptor complex |
| 29 | PTP_med_STAT3_deact | Phosphorylated STAT3 dephosphorylated by PTP3 |
| 30 | PTP_med_STAT1_deact | Phosphorylated STAT1 dephosphorylated by PTP1 |
| 31 | STAT3_SOCS3_ind | Phosphorylated STAT3 inducing SOCS3 protein |
| 32 | STAT3_SOCS1_ind | Phosphorylated STAT3 inducing SOCS1 protein |
| 33 | STAT1_SOCS3_ind | Phosphorylated STAT1 inducing SOCS3 protein |
| 34 | STAT1_SOCS1_ind | Phosphorylated STAT1 inducing SOCS1 protein |
| 35 | IL6R_0 | IL-6R concentration |
| 36 | GP130_0 | GP-130 concentration |
| 37 | IL10R1_0 | IL-10R1 concentration |
| 38 | IL10R2_0 | IL-10R2 concentration |
| 39 | JAK1_0 | JAK1 concentration |
| 40 | JAK2_0 | JAK2 concentration |
| 41 | PTP3_0 | PTP3 concentration |
| 42 | PTP1_0 | PTP1 concentration |
| 43 | SOCS3_degrad | SOCS3 degradation |
| 44 | SOCS1_degrad | SOCS1 degradation |
| 45 | S3_0 | STAT3 concentration |
| 46 | S1_0 | STAT1 concentration |
